# Vocal processing networks in the human and marmoset brain

**DOI:** 10.1101/2024.09.15.613152

**Authors:** Audrey Dureux, Alessandro Zanini, Ravi S. Menon, Stefan Everling

**Author notes:** <u>Corresponding author:</u> Audrey Dureux, Centre for Functional and Metabolic Mapping, Robarts Research Institute, University of Western Ontario, London, Canada.

## Abstract

Understanding the brain circuitry involved in vocal processing across species is crucial for unraveling the evolutionary roots of human communication. While previous research has pinpointed voice-sensitive regions in primates, direct cross-species comparisons using standardized protocols are limited. This study utilizes ultra-high field fMRI to explore vocal processing mechanisms in humans and marmosets. By employing voice-sensitive regions of interest (ROIs) identified via auditory localizers, we analyzed response time courses to species-specific vocalizations and non-vocal sounds using a dynamic auditory-stimulation paradigm. This approach gradually introduced sounds into white noise over 33 seconds. Results revealed that both species have responsive areas in the temporal, frontal, and cingulate cortices, with a distinct preference for vocalizations. Significant differences were found in the response time courses between vocal and non-vocal sounds, with humans displaying faster responses to vocalizations than marmosets. We also identified a shared antero-ventral auditory pathway in both species for vocal processing, originating from the superior temporal gyrus. Conversely, a posterior-dorsal pathway was more prominent in humans, whereas in marmosets, this pathway processed both sound types similarly. This comparative study sheds light on both conserved and divergent auditory pathways in primates, providing new insights into conspecific vocalization processing.

## Introduction

Nonverbal auditory communication, a fundamental mechanism for exchanging socially relevant information across various species, is crucial for discerning the various sounds produced by humans and animals, thereby significantly influencing social interactions ^1–5^.

Recent advances have started to reveal the brain circuits underlying vocal processing in humans and non-human primates. Seminal fMRI studies have identified specific regions within the human temporal lobe that are selectively responsive to vocal sounds as opposed to non-vocal sounds. These include three voice-selective areas along the mid-superior temporal gyrus to the anterior superior temporal gyrus (TVAa, TVAm, TVAp) and associated areas in the premotor and inferior frontal cortex ^6–10^. Further research has identified the superior temporal sulcus (STS) as a pivotal area for processing voice identity and emotional content, with evidence of right hemisphere lateralization ^9,11^.

Parallel investigations in macaques have shown similar vocal processing regions, demonstrating preferential activation of the macaque STS to conspecific vocalizations over other sounds ^12–15^. These findings suggest a hierarchical organization of voice-sensitive regions mirroring those found in humans, indicating an evolutionarily conserved pathway for vocal communication ^16–18^.

Extending these insights to New World marmosets (*Callithrix jacchus*), our prior research has shown that these primates exhibit brain activation patterns in response to vocalizations that align with observations in humans and macaques. We found that marmoset temporal, frontal, and cingulate brain areas exhibit heightened responses to vocalizations compared to non-vocal sounds and scrambled vocalizations ^19,20^, supporting the utility of marmosets as a model for investigating the neural bases of vocal communication ^21,22^.

However, a clearer understanding of the distinct functional roles of each voice-sensitive region across species remains elusive. Until now, no studies have employed comparable experimental paradigms across species to reliably estimate their similarities and differences in this putative voice patch system.

To address this gap, our study investigates the processing of vocalizations in marmosets and humans by employing a dynamic auditory-stimulation fMRI paradigm. This method gradually introduced conspecific vocalizations or non-vocal sounds into an initial white noise sound over a 33-second period, allowing us to explore the temporal dynamics of auditory-responsive regions. By examining the response time courses in multiple voice-sensitive regions of interest (ROIs) identified through fMRI auditory localizers, we were able to compare the neural processing of vocal and non-vocal sounds across species. This approach enabled us to identify both functional similarities and distinctions in the vocal processing pathways of humans and marmosets.

## Results

Nine awake marmoset monkeys and nineteen human subjects participated in a dynamic auditory-stimulation paradigm where conspecific vocalizations and non-vocal sounds were gradually introduced into an initial white noise sound over a 33-second block. Additionally, pure white noise served as a control condition, also presented in 33-second blocks. The primary aim was to determine the response thresholds at which various voice-sensitive ROIs, identified through fMRI auditory localizers, could not only respond to vocalizations but also differentiate between vocal and non-vocal stimuli. To achieve this, we classified the ROIs based on their temporal response patterns—categorized as early, middle, late, or no response—within each species. This classification allowed for a detailed comparative analysis of the functional similarities and differences in auditory vocalization processing between humans and marmosets.

### Auditory localizer and ROI identification in marmosets and humans

We initiated our study by identifying regions responsive to conspecific vocalizations though auditory localizers performed in both marmoset and human subjects. Various contrasts were used, comparing vocal, scrambled vocal, and non-vocal sounds against baseline periods of silence, as well as vocal against scrambled and non-vocal sounds (Figure 1A-E for marmosets and Figure 2A-E for humans). ROIs were delineated exclusively in cortical areas using the NIH marmoset brain atlas ^23^ and the most recent multi-modal cortical parcellation atlas for humans ^24^.

**Figure 1.**
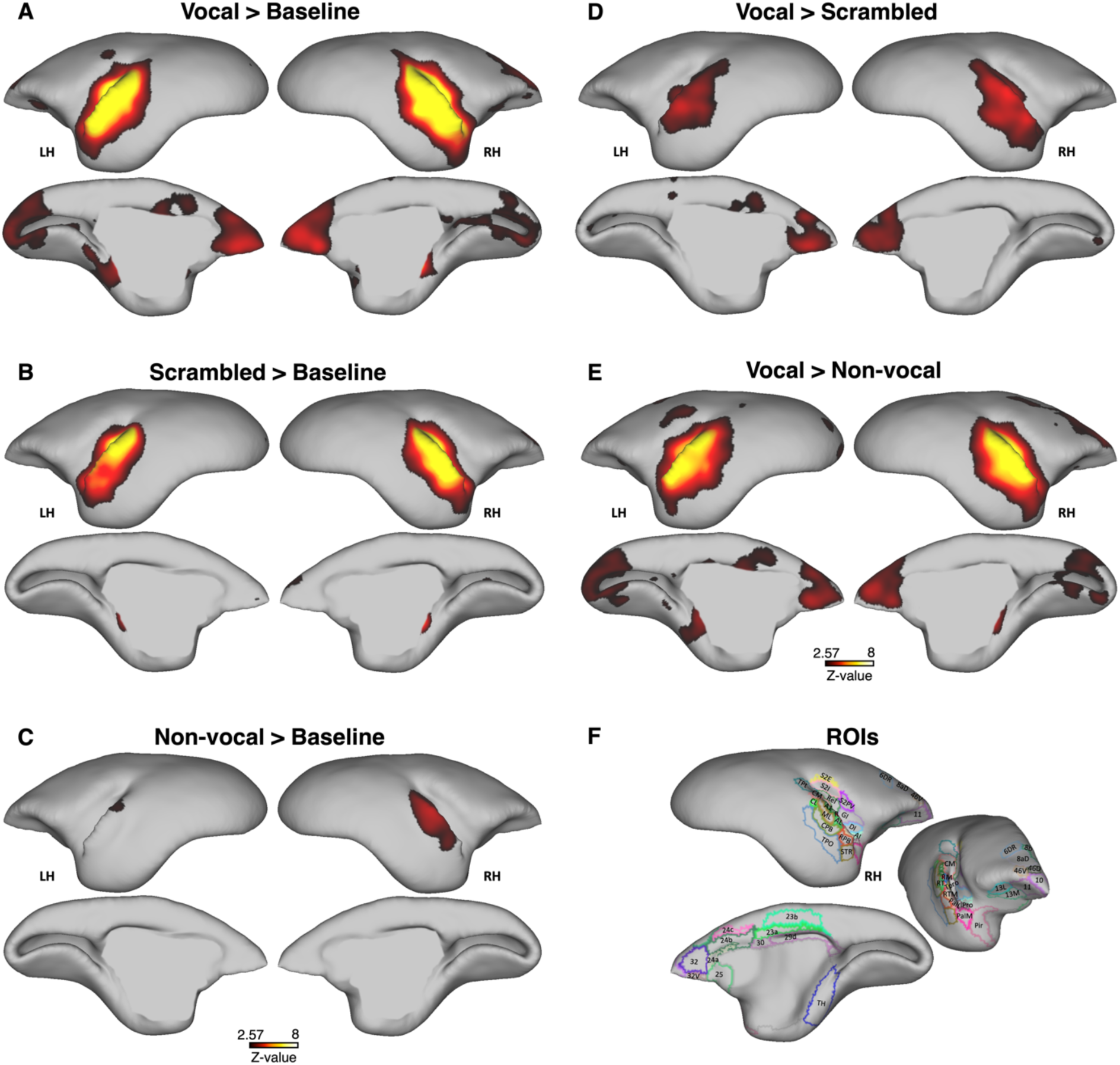
Auditory localizer contrasts and Regions of Interest (ROIs) identification in marmosets. Panels **A-E** depict activation maps for conspecific vocalizations (A), scrambled vocalizations (**B**), non-vocal sounds (**C**) against baseline silence, vocal versus scrambled vocalizations (**D**), and vocal versus non-vocal sounds (**E**), displayed on the left and right fiducial cortical surface. Activation thresholds are set at z-scores > 2.57 (p<0.01, AFNI’s 3dttest++). Panel **F** shows the cortical surfaces with colored outlines delineating the ROIs, based on the NIH marmoset brain atlas ^23^.

**Figure 2.**
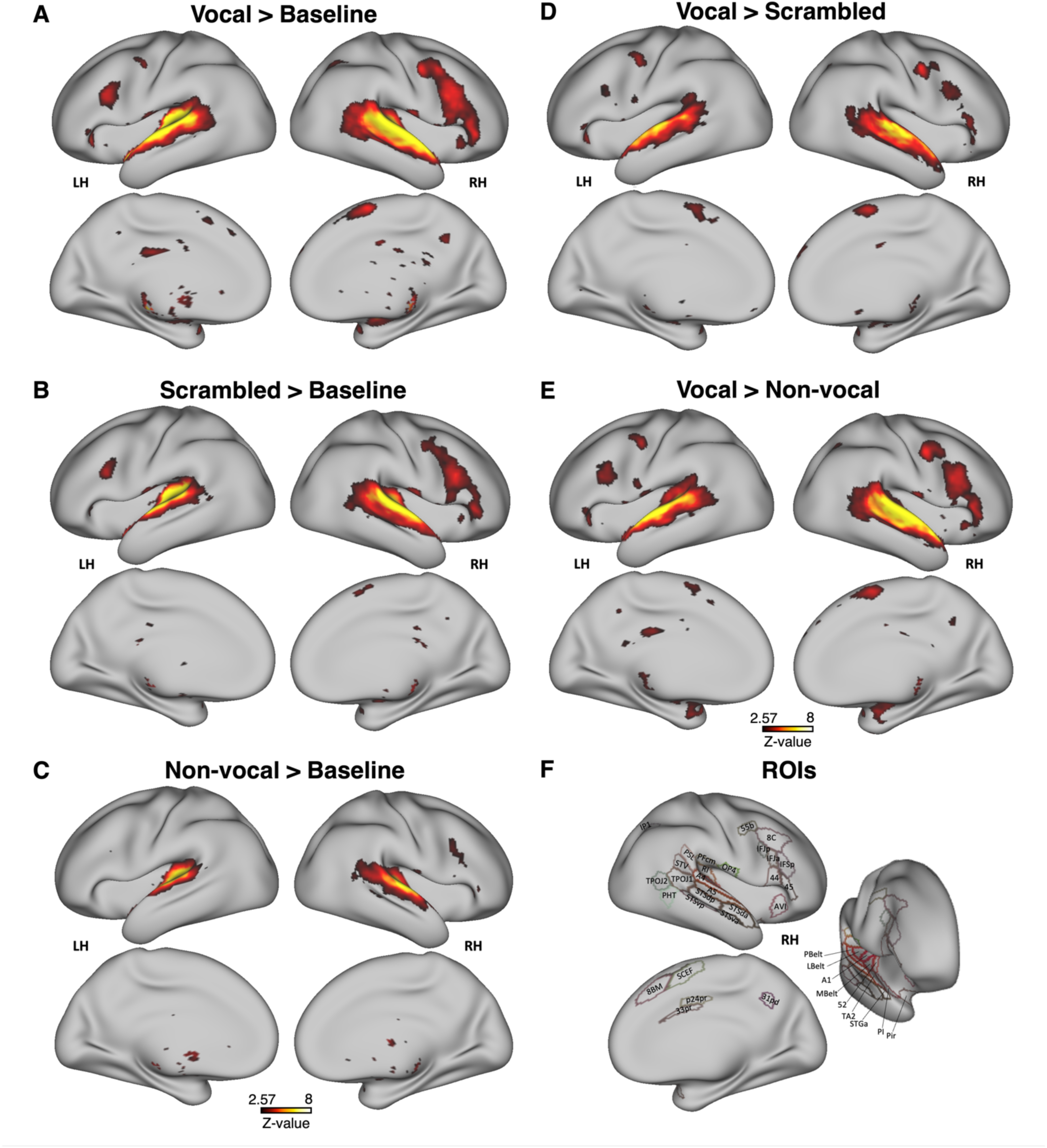
Auditory localizer contrasts and Regions of Interest (ROIs) identification in humans. Panels **A-E** illustrate group activation maps for conspecific vocalizations (**A**), scrambled vocalizations (**B**), non-vocal sounds (**C)** against baseline silence, vocal versus scrambled vocalizations (**D**), and vocal versus non-vocal sounds (**E**), shown on the left and right fiducial cortical surface. Activation thresholds are set at z-scores > 2.57 (p<0.01, AFNI’s 3dttest++). Panel **F** provides the cortical surfaces with colored outlines identifying the ROIs, employing the multi-modal cortical parcellation atlas ^24^.

In marmosets (Figure 1), activations were evident across the auditory core, belt, and parabelt areas, extending into the frontal and cingulate cortex for all contrasts. Notable activations included bilateral responses in core auditory areas such as A1, rostral field (R), and rostral temporal (RT) areas, in belt regions like caudomedial (CM), caudolateral (CL), middle lateral (ML), rostromedial (RM), anterolateral (AL), rostrotemporal medial (RTM) and rostrotemporal lateral (RTL) areas, as well as in parabelt areas like the caudal parabelt (CPB) and rostral parabelt (RPB). Additional activations occurred in several adjacent temporal areas such as the temporo-parietal-occipital area (TPO), superior temporal rostral area (STR), temporoparietal transitional area (TPt), piriform cortex (Pir), and temporal area TH. Secondary somatosensory areas such as external (S2E), internal (S2I), and ventral (S2PV) parts were activated, along with insular regions including dysgranular (DI), granular (GI), agranular (AI) insular areas, insular proisocortex (Ipro), and parainsular areas lateral (PaIL) and medial (PaIM) parts. Significant activations were also observed in prefrontal cortical areas 8b, 8aD, 6DR, 46V and 46D; orbitofrontal areas 10, 11, 13L, 13M; and anterior cingulate areas 24a, 24b, 24c, 25, 32, and 32v, as well as left posterior cingulate areas 23a, 23b, 23c, 29d and 30.

In humans (Figure 2), activations predominantly occurred within the superior temporal gyrus and adjacent insular cortex across all contrasts, including early auditory areas such as A1, 52, and the MedialBelt (MBelt), LateralBelt (LBelt), and ParaBelt (PBelt) Complexes. Additional activations included the retroinsular cortex (RI), para-insular area (PI), and Pir area, alongside auditory association areas A4, A5, and TA2. Posterior temporal cortex activations encompassed areas PFcm, periSylvian language area (PSL), temporo-parieto-occipital TPOJ1 and TPOJ2, PHT, and STV, as well as auditory association areas STSd posterior (STSdp), and STSv posterior (STSvp).

The anterior temporal cortex showed activations in STSv anterior (STSva) and STSd anterior (STSda) areas. Additional activations were also present in lateral prefrontal and inferior frontal areas 55b, 8C, IFJp, IFJa, IFSp, 44, 45, as well as the posterior opercular area (OP4) and the anterior ventral insular area (AVI). Some medial prefrontal and cingulate areas, predominantly in the right hemisphere, were also activated, including 8BM, SCEF, p24pr, 33pr, and 31pd.

In both species, a gradient of responses was observed across the contrasts, from the strongest and most extended activations for vocalizations (Figures 1A and 2A) to the less extensive and weaker activations for non-vocal sounds (Figures 1C and 2C). This gradient led to more pronounced activations when comparing vocal versus non-vocal sounds (Figures 1E and 2E) over vocal versus scrambled sounds (Figures 1D and 2D). The gradient was evident in both the temporal and frontal regions, with prefrontal and cingulate areas being selectively activated by vocal contrasts (Figures 1A, 1D, 1E, 2A, 2D, 2E), with right hemisphere dominance evident in both species.

This analysis delineated 47 cortical ROIs in marmosets and 37 cortical ROIs in humans, as detailed in Figures 1F and 2F respectively.

### Auditory response patterns in marmosets

We analyzed the response time courses for each of the 47 ROIs across 11 time points, specifically focusing on the right hemisphere due to its established lateralization for vocal sound processing in humans ^9,11^ and the higher signal-to-noise ratio provided by our RF coil. The analysis included three conditions: the first involved vocalizations mixed with white noise, varying from 100% white noise with 0% vocalizations to 0% white noise with 100% vocalizations; the second condition followed the same gradient for non-vocal sounds; the third condition was pure white noise, presented consistently across all 11 time points to serve as a control. This design allowed for direct comparisons of auditory processing of vocal and non-vocal sounds compared to pure white noise. Figure 3 shows these three response time courses for each ROI.

**Figure 3.**
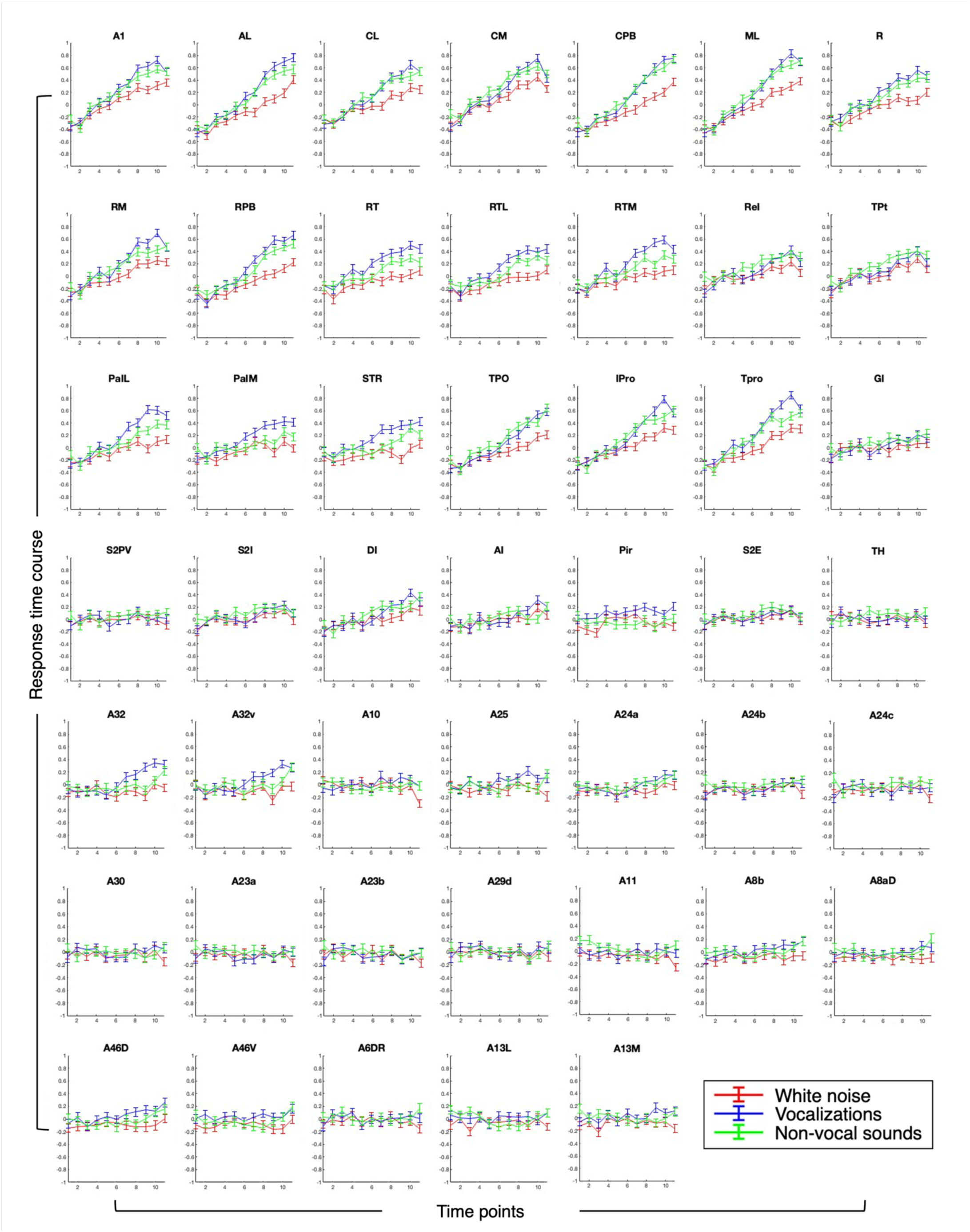
Response time courses of selected ROIs to auditory stimuli in marmosets. The figure illustrates neural responses across 47 selected ROIs in the right hemisphere of marmosets to three auditory conditions: vocalizations (blue) and non-vocal sounds (green) mixed with white noise, and a control condition of pure white noise (red). For vocal and non-vocal sounds, each panel tracks the response from a gradient of 100% white noise at the first time point to 100% of the sound by the eleventh time point, showing a decreasing proportion of white noise and increasing sound intensity. In contrast, the pure white noise control is presented consistently at 100% across all eleven time points to serve as a baseline for comparison. In each plot, the x-axis denotes time points from 1 to 11, and the y-axis indicates the magnitude of neural response, with data points presented as mean responses ± SEM.

Several regions, including A1, AL, CL, CM, CPB, ML, R, RM, RPB, RT, RTL, RTM, ReI, PaIL, PaIM, STR, TPO, IPro, Tpro, 46D, 25, 32, and 32v, showed increasing responses over time to both vocal and non-vocal stimuli, with more pronounced responses to vocalizations, indicative of preferential processing of these biologically relevant sounds. Conversely, parts of the insular and prefrontal cortex (i.e., GI, DI, AI, S2I, S2E, 24a, 8b, 8aD, 46D, 46V, and 13M) as well as TPt area, showed only minimal differentiation between the sound types, suggesting a more generalized role in auditory processing. Areas S2PV, and TH, as well as some prefrontal and cingulate regions, including 24b, 24c, 30, 23a, 23b, 29d, 6DR, 13L, 10, and 11, remained non-responsive across all auditory conditions in this dynamic auditory stimulation paradigm.

Using multiple t-tests at each time point within each ROI, we identified significant points where activations were significantly different for the complex sounds compared to pure white noise. Response timing is visualized in a heat map (Figure 4A), categorizing responses into early (red), middle (orange), and late (yellow) phases. Hierarchical clustering analysis (Figure 4B) delineated the relationships between ROIs under different auditory conditions.

**Figure 4.**
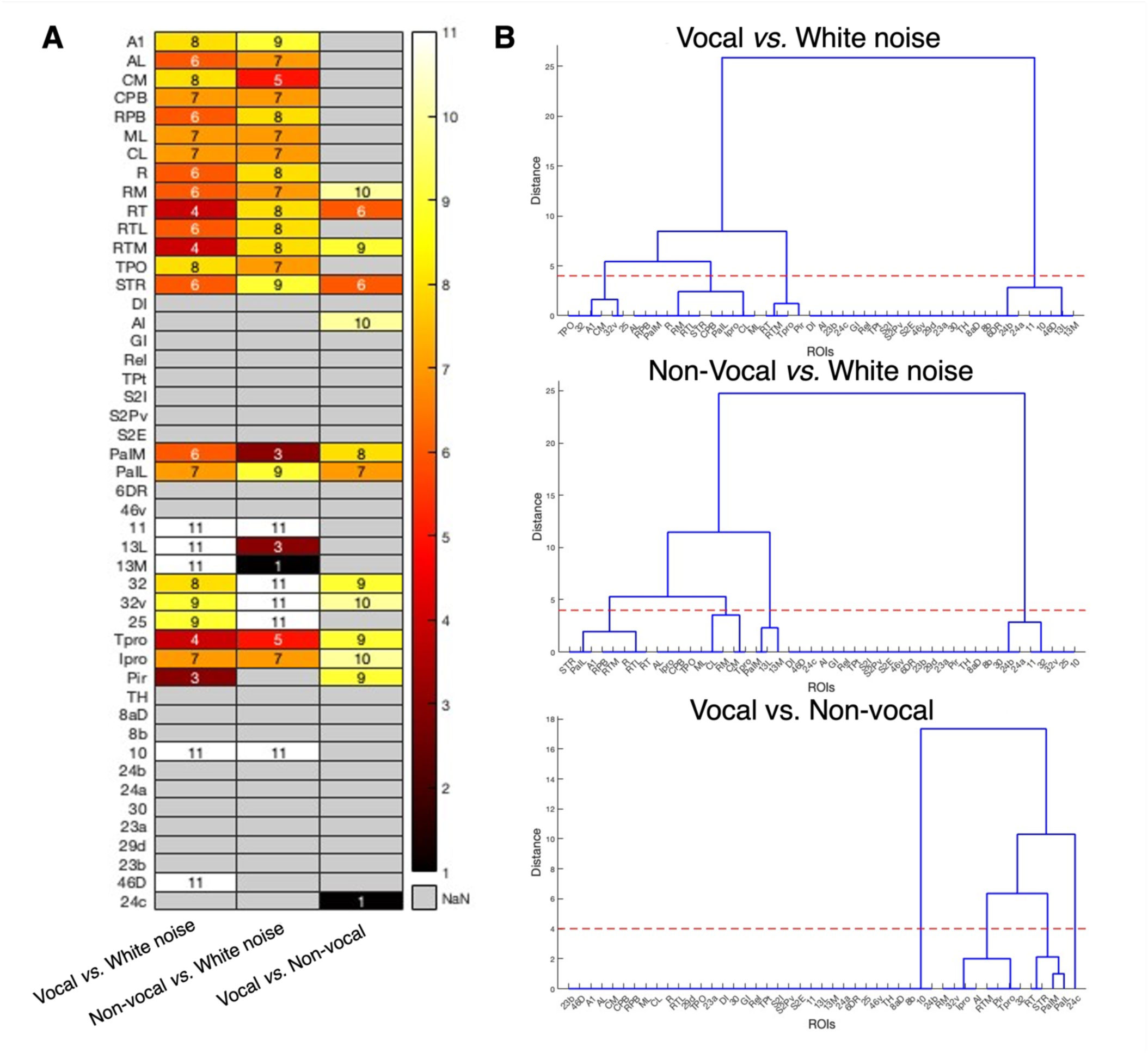
Response onsets and hierarchical clustering of auditory responses in marmosets. **A.** A heat map displays the response onsets of auditory stimuli across 47 right hemisphere ROIs. Each ROI’s response is color-coded to indicate the phase of response: early (red), middle (orange), late (yellow), and no response (grey, represented as NaN). Time points at which significant differences between auditory conditions emerged were identified using two-sided paired t-tests with False Discovery Rate (FDR) post-hoc correction (p < 0.05), revealing specific moments when vocalizations and non-vocal sounds were significantly different from white noise or each other. The x-axis categorizes comparisons, and the y-axis lists the ROIs. **B.** Hierarchical clustering depicts the relationships between ROIs based on their response onsets to three auditory comparisons: vocalizations versus white noise, non-vocal sounds versus white noise, and vocalizations versus non-vocal sounds. Clusters were calculated using Euclidean distance, with linkage distances indicating the degree of functional similarity in response onsets for each comparison.

For vocalizations compared to white noise, temporal regions such as RT, RTM, Tpro, and Pir exhibited early responses, distinguishing between these sounds by the third or fourth time point – equivalent to 20% or 30% vocalization content (Figure 4A). This early detection underscores a heightened sensitivity to vocal sounds and the clustering analysis grouped these areas based on their rapid response profiles (Figure 4B). Middle-phase detection occurred in primary auditory regions (i.e., R, AL, ML, CL, RM, RTL, CPB, RPB), and adjacent anterior temporal areas (i.e., STR, PaIM, PaIL, and Ipro) when vocalizations constituted 50% to 60% of the mix, indicating a more gradual processing capability (Figure 4A). These regions formed a cluster representing moderate response timing (Figure 4B). In contrast, regions like A1, CM, TPO, and anterior cingulate areas 32, 32v, and 25 responded preferentially to vocalizations later in the sequence, requiring higher vocalization content for differentiation (Figure 4A) and were clustered accordingly (Figure 4B). Finally, The largest cluster included prefrontal areas 10, 11, 13L, 13M, and 46D, which responded preferentially to vocalizations only at full intensity (100%), and regions like DI, AI, GI, ReI, TPt, S2I, S2Pv, S2E, 6DR, 46V, TH, 8aD, 8b, 24b, 24a, 30, 23a, 29d, 23b, and 24c, which failed to differentiate at any time point, suggesting a generalized insensitivity or disengagement in vocalization processing in this paradigm (Figure 4A, 4B).

For non-vocal sounds compared to white noise, early responses were present in areas 13M, 13L, and PaIM, although this might be influenced by artifacts at the first time point, corresponding to the initial 100% white noise point. Instead, preferential late-phase responses to non-vocal sounds appear evident in these areas (Figure 4A, also demonstrated in Figure 3). Middle-phase responses occurred in primary auditory areas AL, CM, CPB, ML, CL, RM, and additional temporal regions TPO, Tpro, and Ipro, with significant responses appearing between the fifth and seventh time points, corresponding to 40-60% non-vocal sound, forming a middle response cluster (Figure 4B). Late responses were observed in primary auditory areas A1, RPB, R, RT, RTL, RTM, and anterior temporal area STR, as well as PaIL, occurring at time points eight and nine (Figure 4A), corresponding to 70-80% non-vocal sound. These regions were clustered as late responders (Figure 4B). Very late responses, not apparent until the final time point (100% of the non-vocal sound), occurred in medial prefrontal areas 32, 32v, 25, and lateral prefrontal areas 10, and 11. Finally, several areas including DI, AI, GI, ReI, TPt, S2I, S2Pv, S2E, 6DR, 46V, Pir, TH, 8aD, 8b, 24b, 24a, 30, 23a, 29d, 23b, 46D, and 24c failed to respond higher to non-vocal sounds or exhibited no responses, suggesting a generalized insensitivity or a lack of auditory engagement in this condition, and were regrouped in the same cluster (Figure 4B).

A direct comparison of conspecific vocalizations and non-vocal sounds was also performed to ascertain differences in the processing of these stimuli and further characterize the vocalization responses of specific cortical regions. Among the ROIs, regions such as belt area RT, anterior temporal area STR, and parainsular area PaIL exhibited middle responses, effectively differentiating at 50-60% of the sound mixture. Other areas such as RM, RTM, AI, PaIM, Tpro, Ipro, TH and medial prefrontal areas 32, and 32v, showed late responses, distinguishing between the sound types at higher clarity levels (70-90%) and forming another cluster (Figure 4B). The rest of the areas showed no significant differences between the two sound types (Figure 4A) and were grouped together in the largest cluster (Figure 4B), indicating their generalized auditory response or insensitivity to these auditory conditions.

A comprehensive synthesis of results across auditory comparisons was presented through a dendrogram (Figure 5A), a 3D plot (Figure 5B), and a flat map of the marmoset brain (Figure 5C), illustrating the functional relationships and spatial distribution of ROIs based on their response profiles.

**Figure 5.**
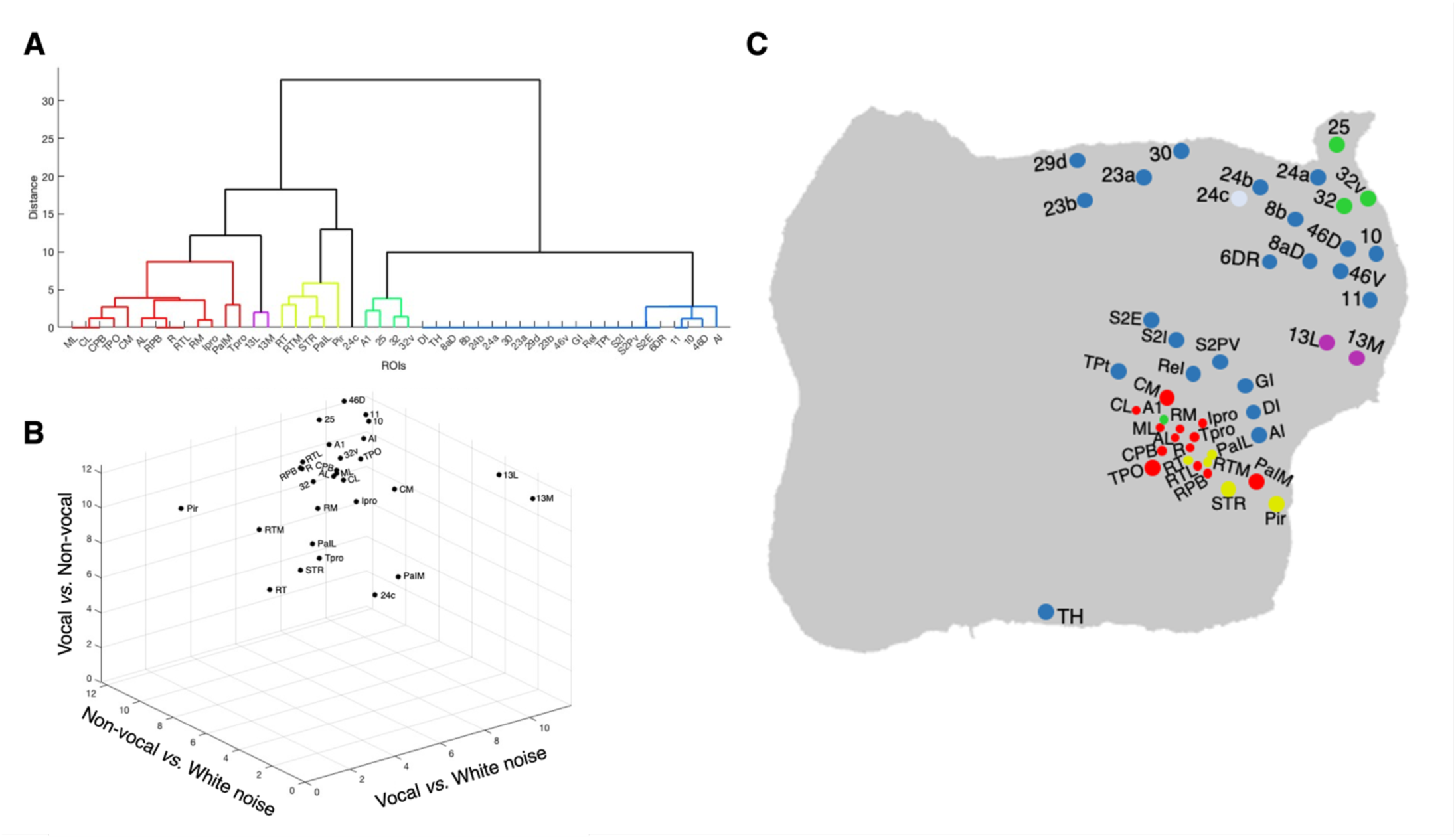
Auditory response comparisons in marmosets. **A.** Dendrogram illustrating the hierarchical clustering of ROIs based on their auditory response profiles to three comparisons: vocalizations versus white noise, non-vocal sounds versus white noise, and vocalizations versus non-vocal sounds. Clusters were derived using Euclidean distance, with linkage distances reflecting the degree of functional similarity in response timings across these comparisons. **B**. A 3D plot depicting the spatial relationships among ROIs according to their functional responses. Axes represent the different auditory comparisons, positioning ROIs to highlight their relative similarities and distinctions based on response times in each comparison. **C.** Flat map of the marmoset brain, color-coded to illustrate each ROI according to the response patterns observed in Panels A and B. This map provides a spatial overview of cortical locations, correlating anatomical positions with functional responses. Colors correspond to clusters identified in Panel A, highlighting the regions’ roles in processing vocal and non-vocal sounds and their ability to differentiate these sounds from white noise and each other.

First, a distinct cluster with high sensitivity to conspecific vocalizations included anterior temporal regions such as RT, RTM, STR, PaIL, and Pir. These regions demonstrated early responses for vocal versus white noise, mid-phase responses for vocal versus non-vocal, and late responses for non-vocal versus white noise. This pattern is highlighted in yellow on both the dendrogram (Figure 5A) and flat map (Figure 5C), with a central position in the 3D plot indicating an important role in vocal sound processing (Figure 5B). Adjacent to these, another cluster encompassed several primary auditory areas such as ML, CL, CPB, TPO, CM, AL, RPB, R, RTL, RM, along with insular areas IPro, PaIM and TPro (Figure 5A), which are anatomically close and presented in red on the flat map (Figure 5C). These regions exhibited similar response profiles for vocal and non-vocal sounds compared to white noise, responding at middle to late phases. They are grouped at the top of the 3D plot (Figure 5B). Notably, some of these regions, like RM, PaIM, TPro and IPro, differentiated between vocal and non-vocal sounds only later in the process. This indicated a general detection capability for auditory stimuli, rather than a rapid response to vocal sounds, although some of these areas showed different responses to these sounds towards the end of the paradigm. A third cluster including the A1 region and anterior cingulate areas 25, 32, and 32, depicted in green, showed predominantly responses to conspecific vocalizations, albeit with a delayed response compared to other temporal regions, suggesting specialized but less sensitivity to vocal sounds (Figure 5A, Figure 5B and Figure 5C). Orbitofrontal areas 13L and 13M, depicted in pink, were differentiated from others due to their unique response patterns to non-vocal sounds, likely influenced by an artifact at the first time point (see above). Similarly, the cingulate area 24c is also separated from the other regions. Finally, the largest cluster comprised peripheral areas like the temporal TH region, along with various prefrontal and posterior cingulate areas, shown in blue, which exhibited minimal or no responses across all conditions (Figure 5A and Figure 5C). This cluster is positioned away from the core operational axes in the 3D plot (Figure 5B), indicating its limited role in auditory processing.

### Auditory response patterns in humans

Following the methodology applied to marmosets, we analyzed auditory responses in humans across 11 time points. We focused on the right hemisphere, not only because of its established lateralization for vocal sound processing ^9,11^ but also due to the higher signal-to-noise ratio provided by our coil in this hemisphere ^25^. Figure 6 presents the response time courses, illustrating how the 37 ROIs responded to vocal and non-vocal stimuli relative to the white noise condition.

**Figure 6.**
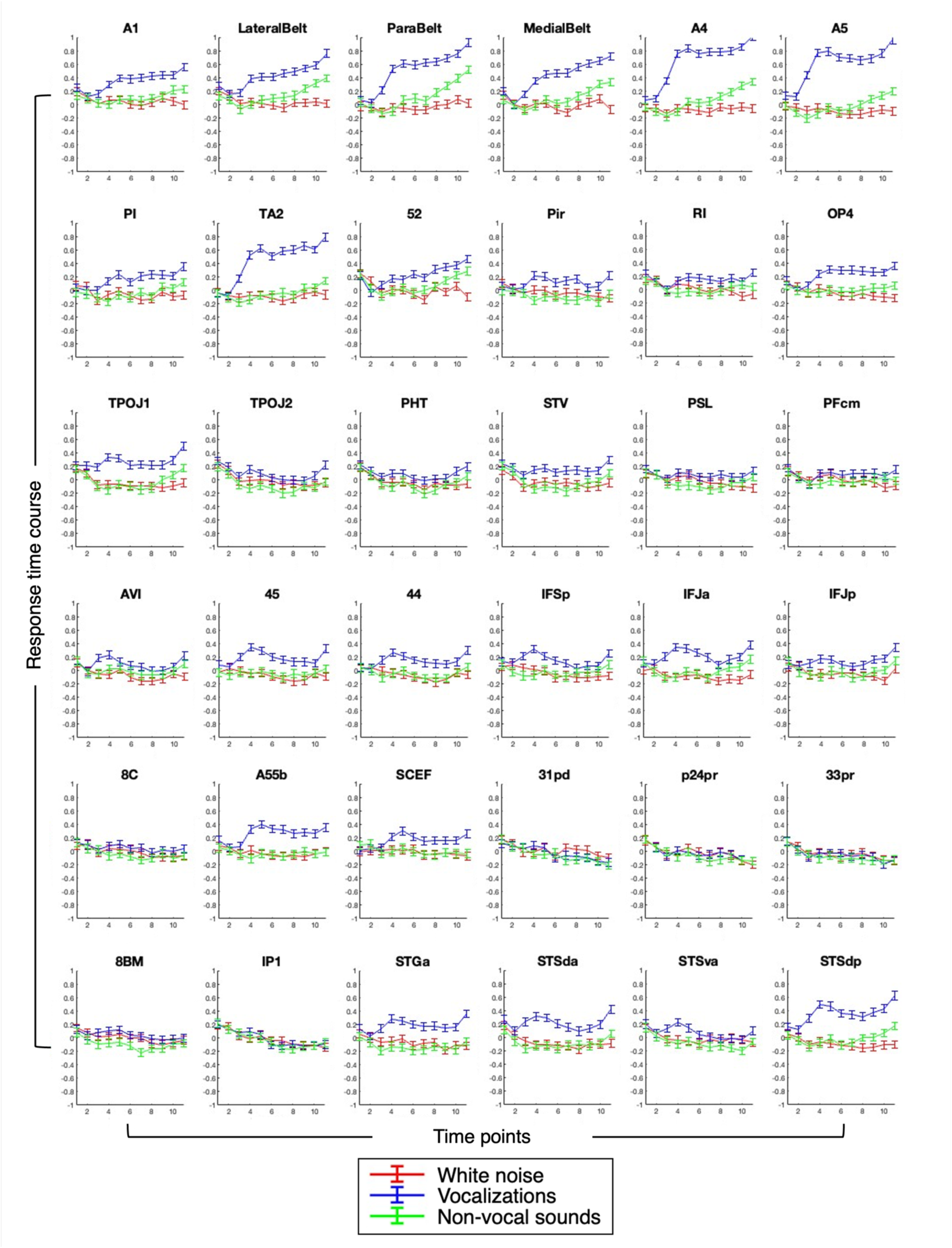
Response time courses of selected ROIs to auditory stimuli in humans. This figure displays the neural activity within 37 selected ROIs of the right hemisphere in response to three types of auditory stimuli: white noise (red), vocalizations (blue), and non-vocal sounds (green), plotted over 11 time points (33s). Each time point transitions from 100% white noise to 100% of the specific sound, with the control condition of pure white noise maintained at 100% across all time points for baseline comparison. Each panel charts the magnitude of neural responses, marked on the y-axis, against the time points on the x-axis, with data points presented as mean responses ± SEM.

Several regions, including A1, LBelt, PBelt, MBelt, A4, A5, area 52, TPOJ1, IFSp, IFJa, IFJp, and STSdp, displayed increasing responses to both vocalizations and non-vocal sounds, with notably earlier and more pronounced responses to vocalizations, suggesting heightened sensitivity to these socially relevant sounds. Notably, areas such as PI, TA2, OP4, 45, 44, 55b, SCEF, STGa, and STSda responded almost exclusively to vocalizations, showing minimal or no response to non-vocal sounds and pure white noise. Conversely, RI, Pir, TPOJ2, PHT, STV, AVI, and STSva, demonstrated very weak responses to vocalizations. Additional areas, including PSL, PFcm, IP1, and medial and lateral prefrontal areas 8C, 31pd, p24pr, 33pr, and 8BM, remained unresponsive to all tested stimuli.

We conducted multiple t-tests at each time point within each ROI to identify significant differences in neural responses between conditions, distinguishing responses to pure white noise from those elicited by vocalizations and non-vocal sounds, and between vocalizations and non-vocal sounds. Results are visualized in a heat map (Figure 7A) and separate dendrograms (Figure 7B) to illustrate the response onsets and establish the functional relationships among ROIs.

**Figure 7.**
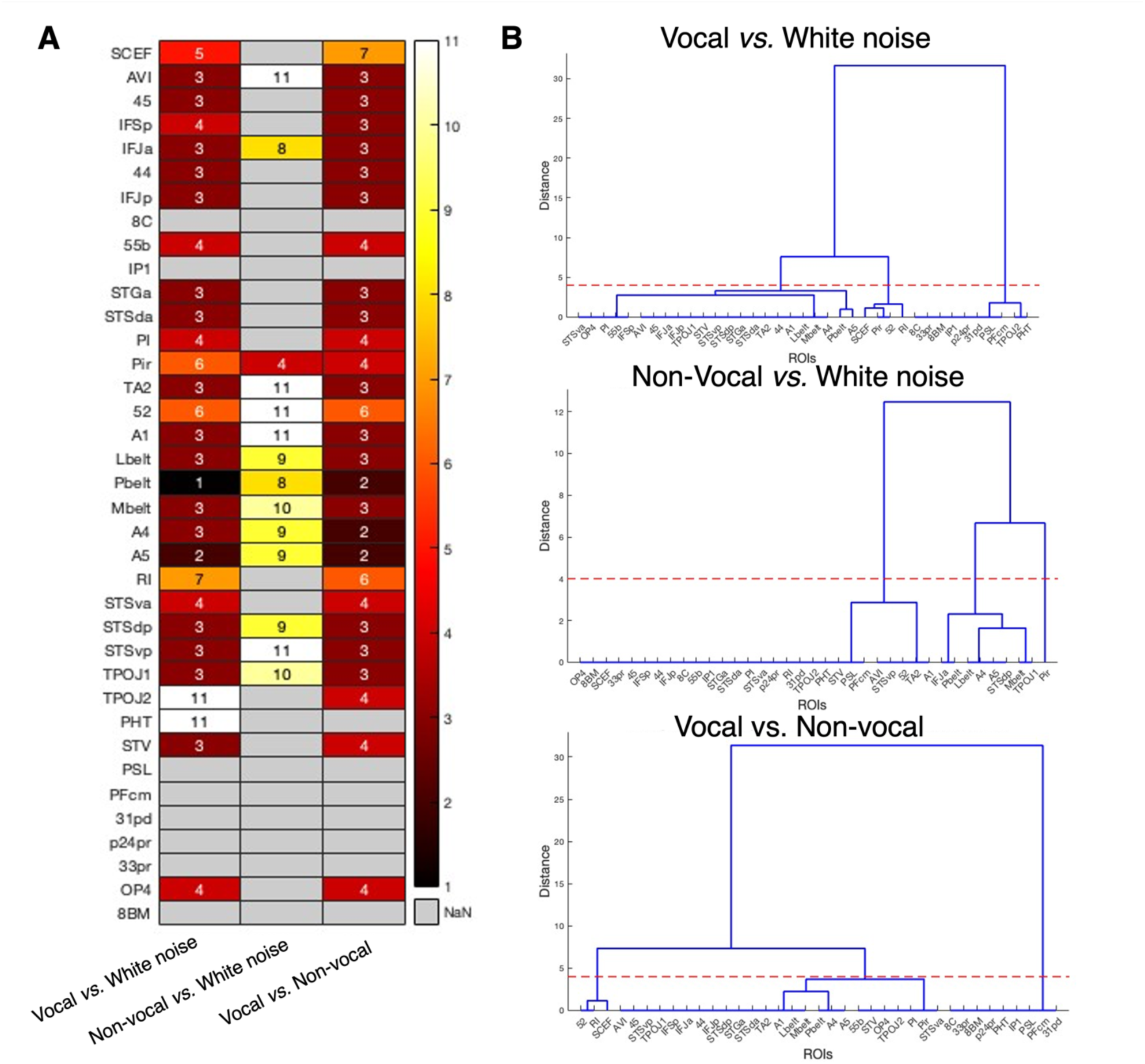
Response onsets and hierarchical clustering of auditory responses in humans. **A.** The heat map illustrates the significant response onsets for auditory stimuli across various ROIs, color-coded to represent early (red), middle (orange), and late (yellow) responses, with non-responsive regions in grey. Significance was determined using two-sided paired t-tests with False Discovery Rate (FDR) correction (p < 0.05), comparing vocalizations to white noise, non-vocal sounds to white noise, and vocalizations to non-vocal sounds. **B.** The dendrogram displays the hierarchical clustering of ROIs based on their response onsets to the three auditory comparisons. Clusters are formed using Euclidean distance, with linkage distances indicating the degree of functional similarity in response onsets for each comparison.

For the comparison between vocalizations and white noise, several regions demonstrated early responses. Inferior frontal regions such as areas 44, 45, IFJa, IFJp, and the premotor area 55b, as well as early auditory areas A1 and various belt complexes located in the superior temporal gyrus along with the auditory association areas STGa, STSda situated in the anterior temporal cortex and adjacent insular areas AVI and PI, responded quickly to vocalizations, responding selectively to these sounds by the third or fourth time point, which corresponded to 20% to 30% vocalization content (Figure 7A). Notably, auditory area A5 responded to vocalizations even earlier with only 10% of sound. These regions were collectively grouped in the first cluster of the dendrogram (Figure 7B), reflecting their fast response profile and heightened sensitivity to vocal sounds. Other areas, including cingulate area SCEF, the inferior temporal gyrus areas 52, and RI, as well as the insular area Pir, responded to vocalizations during a middle phase, when they constituted between 40% and 60% of the auditory mix (Figure 7A). These regions formed another cluster, indicating a moderate response time suggesting a gradual processing capability of these regions (Figure 7B). Lastly, several regions either failed to respond to vocal sounds at any intensity (i.e., 8C, IP1, PSL, PFcm, 31pd, p24pr, 33pr, 8BM), or only did so when vocalization content reached 100% (i.e., posterior temporal areas TPOJ2 and PHT), forming the last cluster in the dendrogram (Figure 7A and 7B).

For the comparison between non-vocal sounds and white noise, most regions, including the inferior frontal area IFJa, along with early auditory areas LBelt and PBelt, and auditory association areas A4, A5, and STSdp, responded to non-vocal stimuli relatively late, between 70% to 80% non-vocal sound content (Figure 7A). Additionally, several regions responded to non-vocal sounds only at 90-100% non-vocal content. These regions included early auditory areas A1 and 52, the MBelt complex, auditory association area TA2, and posterior temporal areas such as STSvp and TPOJ1, as well as the adjacent insular area AVI (Figure 7A). An exception was the Pir region, which responded to non-vocal sound at 30% of the sound presence. All other regions did not differentiate between the two stimuli at any time points (Figure 7A). The response patterns across these regions led to the formation of three distinct clusters within the dendrogram (Figure 7B), representing the varying response timings (Figure 7A).

A direct comparison of vocal and non-vocal sounds revealed that several regions, particularly those in the superior temporal gyrus (TA2, A1, LBelt, PBelt, MBelt, STGa), anterior and posterior temporal lobes (STSda, STSva, STSdp, STSvp, TPOJ1, TPOJ2, STV), and adjacent insular area (AVI, PI, Pir), along with regions in the prefrontal and inferior frontal cortex (55b, 44, 45, IFSp, IFJa, IFJp, OP4), exhibited higher sensitivity to conspecific vocalizations compared to non-vocal sounds, with a significant difference in response times occurring between 10% and 30% sound mixed with 70% to 90% white noise (Figure 7A). These regions formed the largest cluster in the dendrogram (Figure 7B). A second cluster, including the cingulate area SCEF and the early auditory areas 52 and RI, demonstrated moderate sensitivity, with notable differences in responses appearing at 50% and 60% of the vocal sound (Figures 7A and 7B). The final cluster consisted of posterior temporal regions such as PHT, PSL, PFcm, the parietal area IP1, the prefrontal area 8C, and the medial prefrontal areas 31pd, p24pr, 33pr, and 8BM, which showed no significant differences between the two sounds or were unresponsive (Figures 7A and 7B).

A comprehensive synthesis of auditory comparisons is represented through a dendrogram (Figure 8A), a 3D plot (Figure 8B), and a flat map of the human brain (Figure 8C).

**Figure 8.**
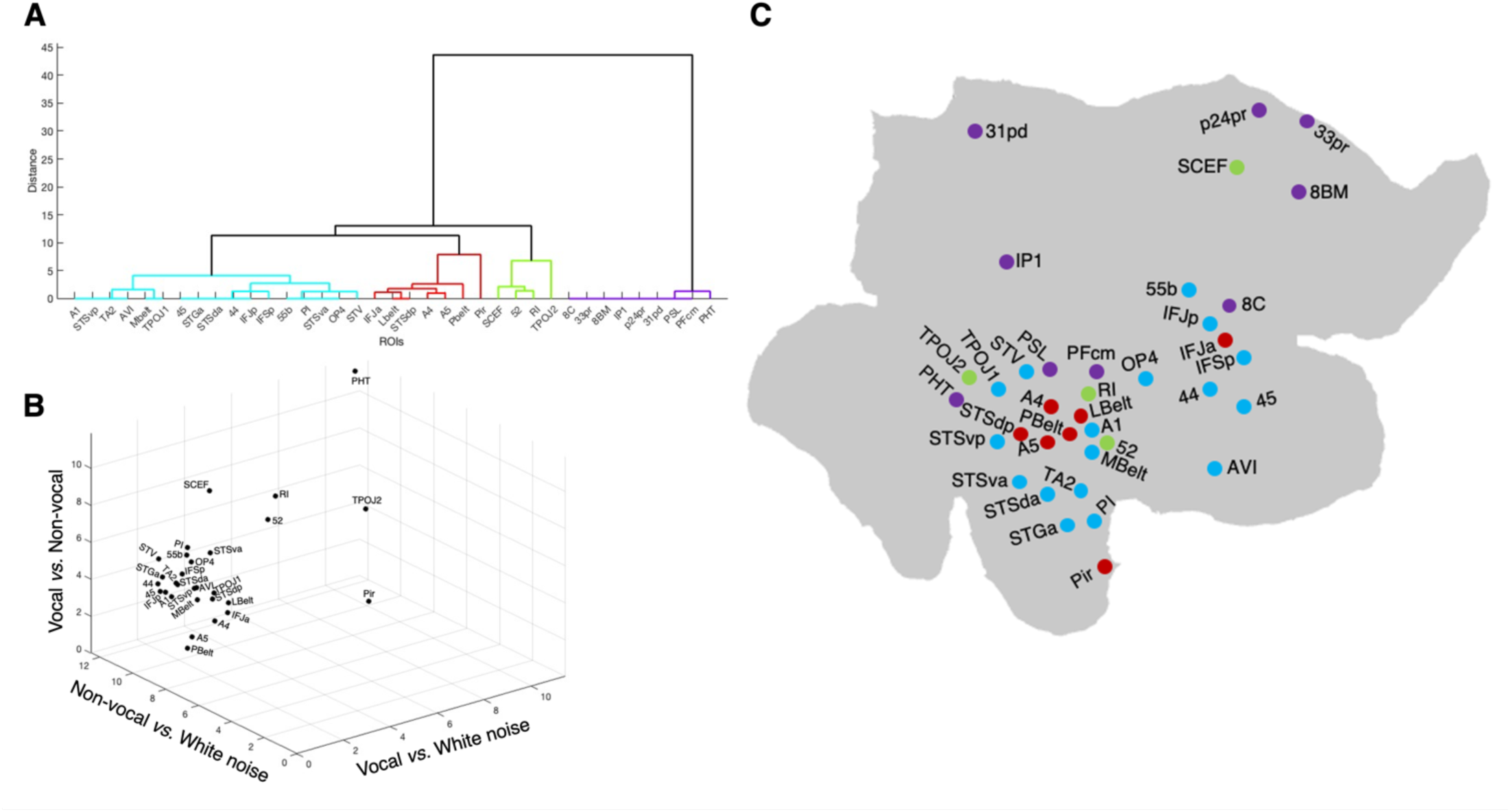
Auditory responses across comparisons in humans. **A.** Dendrogram illustrating the hierarchical clustering of ROIs based on their response profiles to three auditory comparisons: vocal versus white noise, non-vocal versus white noise, and vocal versus non-vocal sounds. Clusters are formed using Euclidean distances to highlight functional similarities among ROIs. **B.** A 3D plot delineates the spatial and functional relationships among ROIs, with axes corresponding to the different auditory comparisons, demonstrating how regions align or differ in their response patterns. **C.** A flat map of the human brain displays each ROI’s anatomical location, color-coded according to the response patterns and clusters identified in Panel A, showcasing the regions’ abilities to differentiate between vocal and non-vocal sounds, as well as their responses of these sounds relative to white noise.

The largest cluster identified in the dendrogram included regions with early responses for vocalizations compared to both white noise and non-vocal sounds, highlighting a pronounced sensitivity to vocal stimuli. Within this cluster, in light blue, two distinct subgroups were notable (Figure 8A and Figure 8C). The first subgroup included superior temporal areas TA2, A1, MBelt, posterior temporal areas STSdp, TPOJ1, and the insular area AVI. These regions only responded to non-vocal sounds at 100%. The second subgroup consisted of anterior temporal areas STGa, STSda, STSva, the posterior temporal area STV, the insular area PI, and frontal areas 44, 45, IFJp, IFSp, 55b, OP4. These areas showed no differentiation between non-vocal sounds and white noise. A second cluster, highlighted in dark red, included auditory areas A4, A5, PBelt, and LBelt located in the superior temporal gyrus, along with the posterior temporal area STSdp, the insular area Pir and the inferior frontal area IFJa. These regions responded to both vocal and non-vocal sounds with a stronger response to vocal stimuli (Figure 8A and Figure 8C). The regions of these two clusters were closely positioned in the 3D plot, near the origin along the axes comparing non-vocal sounds versus white noise and vocal versus non-vocal sounds, indicating their sensitivity to vocalizations (Figure 8B).

The third cluster included more dispersed regions such as TPOJ2, the early auditory areas 52 and RI in the STS and the cingulate area SCEF, colored in light green. These regions showed later response onsets for both vocalizations versus white noise and vocalizations versus non-vocal sounds, without differentiating non-vocal sounds from white noise (Figure 8A and Figure 8C). This pattern indicates a primary but delayed focus on vocal sound processing, distinctly set apart in the 3D plot (Figure 8B).

Lastly, a cluster in purple included regions such as 8C, 33pr, 8BM, IP1, p24pr, 31pd, PSL, PFcm, and PHT, showing no responses to auditory stimuli (Figure 8A and Figure 8C). These regions were positioned away from the main axes in the 3D plot, indicating minimal involvement in the dynamic auditory-stimulation paradigm (Figure 8B).

### Comparison of responses between humans and marmosets

Our analysis of auditory responses between humans and marmosets revealed both similarities and distinct differences in how each species processes auditory stimuli. While both species demonstrated the ability to detect conspecific vocalizations earlier than non-vocal sounds, humans exhibited earlier responses to vocalizations, typically between 20-30% vocalization content relative to white noise. In contrast, marmosets generally required 40-50% vocalization content for similar response onsets. Notably, several primary auditory cortex areas and adjacent insular regions, responded to non-vocal sounds earlier relative to white noise in marmosets than in humans. For example, marmoset areas such as AL, CM, ML, CL, RM, CPB, and adjacent areas like TPO, Tpro, and Ipro, responded to these sounds between 40-60% or later at 70-80% for primary auditory regions like A1, R, RT, RPB, RTL, RTM, and adjacent STR and PaIL. Conversely, human auditory regions including A4, A5, LBelt, PBelt, STSdp, and the inferior frontal area IFJa responded to non-vocal sounds much later, typically between 70-90%, with some superior temporal gyrus areas (A1, TA2, 52, MBelt) and posterior temporal areas (TPOJ1, STSvp) only responding at 90-100%.

Furthermore, nearly all responding human auditory regions showed different responses between vocal and non-vocal sounds at lower mixing ratios (10-30%), while marmoset regions typically required at least 50% sound presence, such as RT, STR, and PaIL, or even higher percentages (70-90%) for regions like RM, RTM, AI, PaIM, TH, Ipro, Tpro, 32, and 32v. In humans, only a few regions like SCEF, 52, and RI exhibited such delayed responses. Several marmoset regions did not show any response differences between the two sounds, in contrast to only a few regions in humans, suggesting a higher sensitivity for conspecific vocalizations in humans. These regions included primary auditory areas A1, AL, ML, CL, CM, R, RTL, CPB, RPB, adjacent TPO, and prefrontal areas 10, 11, 13L, and 13M.

Figure 9 illustrates this cross-species auditory processing comparison. In figure 9A, the ROIs are categorized into distinct clusters based on their ability to respond differentially to vocalizations and white noise, non-vocal sounds and white noise, and vocalizations versus non-vocal sounds, without consideration of response onsets (early, middle, or late). Figure 9B integrates these findings with anatomical representations, linking the spatial distribution of ROIs with their functional clustering, color-coded according to their groupings in the dendrogram.

**Figure 9.**
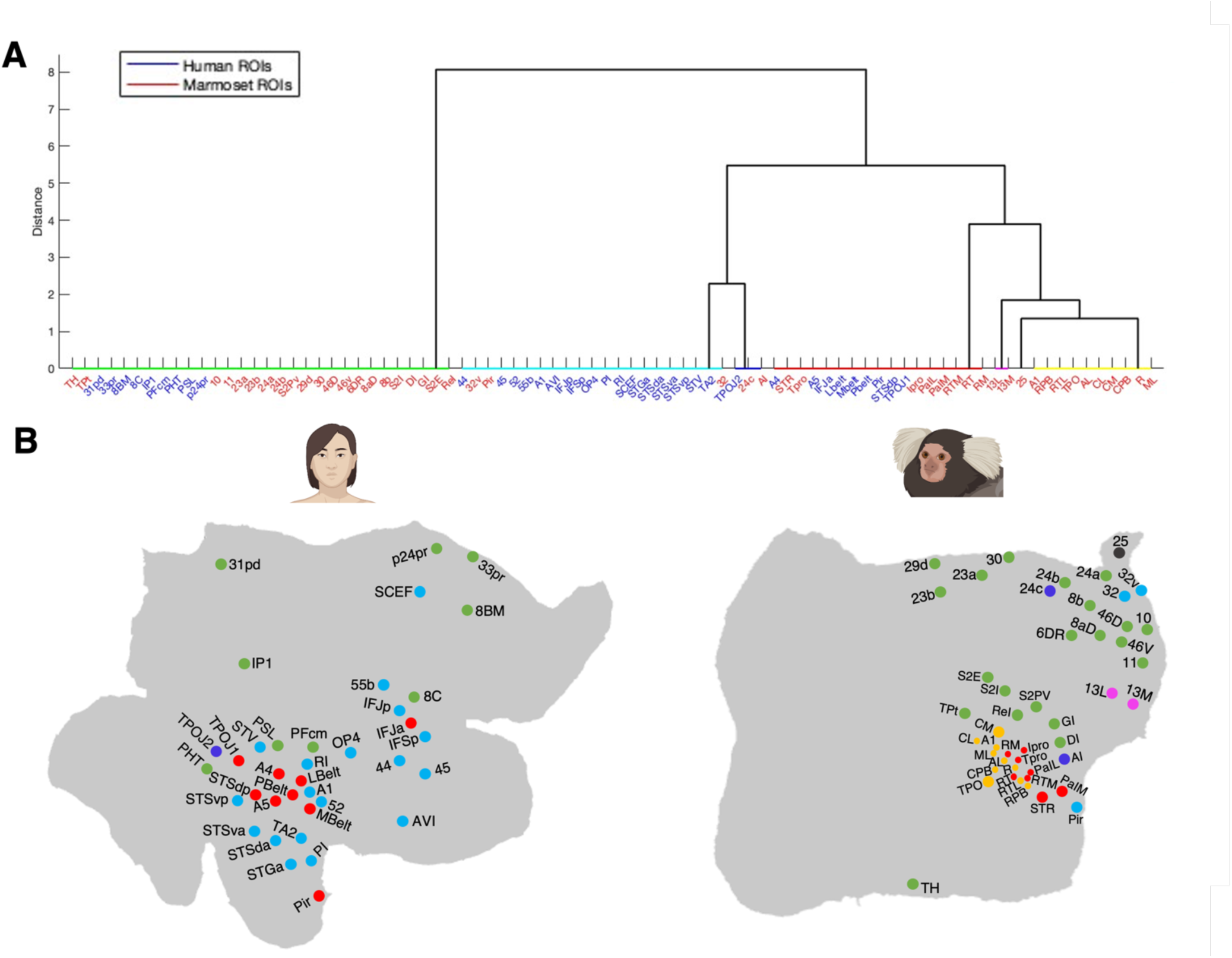
Comparative analysis of auditory processing between humans and marmosets. **A.** Dendrogram presenting the hierarchical clustering of ROIs from both humans and marmosets, illustrating similarities and differences in auditory processing between the two species. ROIs are grouped based on their response patterns to different sounds—distinguishing between vocalizations and white noise, non-vocal sounds and white noise, and vocal and non-vocal sounds—without consideration of response onsets. Clustering is shown with human ROIs in blue and marmoset ROIs in red, calculated using Euclidean distance. **B.** Anatomical representations on flat maps of both human and marmoset brains showing the spatial distribution of these ROIs. Each dot represents an ROI, color-coded according to the cluster it belongs to in the dendrogram (Panel A). This visualization provides a clear correlation of anatomical location with functional response patterns across species, highlighting regions with similar response profiles.

A prominent cluster, depicted in light blue, encompassed regions that significantly distinguished between vocalizations and both white noise and non-vocal sounds. However, these regions did not differentiate between non-vocal sounds and white noise, or they only did so when the non-vocal sound presence reached 100%. This cluster included human prefrontal areas 44, 45, IFJp, IFSp, OP4, and the premotor area 55b, along with areas in the superior temporal gyrus such as early auditory areas A1, 52, RI, and auditory association areas TA2 and STGa. It also comprised anterior temporal areas STSda, STSva, posterior temporal areas STSvp, STV, adjacent insular areas AVI, PI, and cingulate area SCEF, alongside marmoset cingulate areas 32 and 32v, and the anterior and ventral temporal area Pir. Isolated in grey, marmoset area 25 formed its own cluster due to its unique response to vocalizations over white noise. Despite potentially being grouped with the light blue cluster due to its ability to respond to vocalizations, its lack of significant differentiation between vocal and non-vocal sounds sets it apart.

Another small cluster in dark blue, containing human TPOJ2 and marmoset areas 24c and AI, exhibited significant differentiation solely between vocalizations and non-vocal sounds. This pattern suggests a uniform processing of both sound types throughout the auditory sequence, with differentiation only becoming apparent towards the end.

A robust red cluster showed strong responses across all auditory comparisons. In humans, it included regions within the superior temporal gyrus such as the early auditory areas LBelt, MBelt, PBelt, and auditory association areas A4, A5, the anterior ventral temporal area Pir, posterior temporal area STSdp, TPOJ1, and the inferior frontal area IFJa. In marmosets, this cluster featured primary auditory areas in the superior temporal gyrus (core RT, belt RM, RTM), the Tpro area, anterior temporal area STR, and adjacent insular areas Ipro, PaIL, and PaIM. These regions were central in responding to both vocal and non-vocal auditory stimuli and could differentiate between them. Additionally, a small fifth cluster, shown in pink, included marmoset orbitofrontal areas 13L and 13M, characterized by their unusual responses to non-vocal sounds at the initial time point. A unique yellow cluster to marmoset regions regrouped those that showed higher responses for vocalizations than white noise and for non-vocal sounds versus white noise, yet did not show differences between vocal and non-vocal sounds. These regions were primarily located in the superior temporal gyrus, including primary auditory areas A1, R, RTL, AL, CL, CM, ML, RPB, and CPB, as well as the adjacent TPO region. Finally, a green cluster included regions in both species that had no significant selectivity, indicating either generalized auditory processing or insensitivity to the conditions tested. In humans, these areas were predominantly located in peripheral regions of the lateral and medial prefrontal cortex, such as 8C, 31pd, 33pr, p24pr, and 8BM, as well as the inferior parietal area IP1, and posterior temporal areas PFcm, PSL, and PHT. In marmosets, similar non-selectivity was observed in peripheral cortical areas including the lateral and medial prefrontal cortex, such as 10, 11, 8b, 8aD, 46V, 23a, 23b, 24a, 24b, 29d, 30, the premotor area 6DR, and extend to some areas of the insular cortex (i.e., DI, GI, ReI), the posterior temporal area TPt, the medial temporal area TH, and the secondary somatosensory cortex (i.e., S2I, S2E, S2PV). Prefrontal area 46D, which selectively responded to vocalizations compared to white noise only at full sound presence, was included in this cluster.

## Discussion

We explored and compared vocalization processing mechanisms in humans and marmosets using ultra-high field fMRI by focusing on multiple voice-sensitive regions of interest (ROIs) identified through auditory localizers. These ROIs corresponded to previously identified regions involved in vocalization processing in both species, as highlighted in foundational studies ^6–10,19,20^.

We evaluated the processing of species-specific vocalizations and non-vocal sounds using a dynamic auditory-stimulation paradigm that gradually introduced these distinct sounds into an initial white noise stimulus. This approach allowed us to uncover the auditory processing capabilities of each region, as well as the specific sound-noise combinations that elicited responses. Our findings revealed patterns of both generalized auditory responses and distinct activations for conspecific vocalizations across various brain regions in both species.

In marmosets, a wide array of auditory cortex areas across the core, belt, and parabelt within the superior temporal gyrus (i.e., A1, AL, R, RT, RM, ML, RTL, RTM, TPro), lateral and posterior temporal cortex (i.e., CL, CM) and anterior temporal cortex (RPB, STR), along with regions in the lateral and inferior temporal cortex (i.e., CPB, TPO), insula (i.e., PaIM, PaIL, IPro), and dorsolateral prefrontal and orbitofrontal cortex (46D, 13M), demonstrated strong responses to both vocalizations and non-vocal sounds. However, these regions showed varying response gradients, with most auditory areas identifying conspecific vocalizations from white noise earlier than non-vocal sounds, often identified at early stages (10 to 30%) or mid-stages (40 to 60%), contrasting with later responses (70-90%) for non-vocal sounds. This distinct pattern was especially pronounced in regions within the superior temporal gyrus like R, RT, RTM, RTM and PaIL, as well as in anterior temporal cortex in RPB and STR and in prefrontal area 46D. Notably, anterior medial prefrontal areas 32, 32v, and 25 specifically responded to vocalizations and required the complete absence of white noise to discern non-vocal sounds, only when those reached 100% of sound presence. Certain areas, including superior temporal areas RT, RM, RTM, TPro and anterior temporal area STR, alongside ventral temporal area Pir and adjacent insular areas PaIM, PaIL, IPro, as well as anterior cingulate areas 32 and 32v, showed significant difference in responses for vocalizations compared to non-vocal sounds, highlighting a specialized neural circuit that prioritizes biologically relevant vocal sounds crucial for communication and social interactions.

Humans exhibited similar response patterns within the superior temporal gyrus (i.e., auditory areas A1, 52, MBelt, PBelt, LBelt and auditory association areas A4, A5, TA2), posterior temporal areas (i.e., STSdp, STSvp, and TPOJ1), and adjacent ventral temporal areas (i.e., AVI, Pir), and the inferior frontal area IFJa. These regions responded to conspecific vocalizations from early to mid-stages, contrasting with the much later identification of non-vocal sounds (70-90%, or requiring 100% sound purity). Furthermore, numerous human regions, including the frontal (44, 45, IFSp, IFJp), premotor (55b), insula (PI), posterior temporal (TPOJ2 and STV), and anterior temporal cortex (STSGa, STSda, STSva), as well as the superior temporal RI area and the cingulate area SCEF, were distinctly capable of differentiating between conspecific vocalizations and white noise and between vocalizations and non-vocal sounds. These observations highlight the enhanced sensitivity or specialized processing capabilities of these regions for conspecific vocalizations. This finding aligns with prior research suggesting that during the perception of vocal signals, these signals traverse the ascending auditory pathway, ultimately engaging both low-level (A1) and high-level auditory cortex (superior temporal sulcus). From the superior temporal sulcus (STS), two distinct streams emerge: an antero-ventral stream originating from the anterior STS, facilitating the identification of diverse vocal information, including the sender’s identity (temporal pole) and referential meaning (lateral prefrontal cortex), extending to the anterior cingulate cortex, and a postero-dorsal stream from the posterior STS involved in encoding auditory-motor information, including temporal information (premotor cortex) ^1,26^. Electrophysiological studies, such as those by De Lucia et al. (2010), have shown that responses to human versus animal vocalizations begin markedly faster, from 169–219 ms post-stimulus onset within the right superior temporal gyrus, with subsequent differentiation around 291–357 ms within the inferior prefrontal gyrus ^27^. Capilla et al. (2012) further identified an early voice-preferential response peaking at approximately 220 ms, distinguishing between vocal and non-vocal sounds as soon as 150 ms after stimulus onset, localized along the mid-STS/STG ^28^. Together with these studies, our findings highlight early, voice-selective cerebral mechanisms with a response time course that appears comparable to face processing ^29,30^, extending from the superior temporal gyrus to the anterior temporal cortex and the lateral prefrontal cortex, and ending in the cingulate area SCEF, which exhibits later vocalization responses compared to other regions.

Our finding show that both humans and marmosets not only possess similar voice-selective regions but that these regions also exhibit a gradient of vocal selectivity from core auditory temporal areas to the prefrontal cortex. However, the response time courses of voice-selective areas exhibited both similarities and differences between humans and marmosets.

Concerning similarities, firstly, several superior temporal areas, recognized as key auditory regions, along with adjacent anterior and posterior temporal areas in both species, responded to both conspecific vocalizations and non-vocal sounds compared to white noise, with earlier response times for vocalizations than for non-vocal sounds. In marmosets, these included core (R, RT), belt (RTL, RTM), and parabelt (RPB) auditory regions, PaIL area, and anterior temporal areas STR. In humans, these encompassed the superior temporal gyrus (A1, A4, A5, MBelt, PBelt, LBelt, TA2, 52), including core, belt and parabelt auditory regions, and adjacent insula AVI and Pir areas, as well as posterior temporal areas (STSdp, STSvp, TPOJ1) and the inferior frontal area IFJa.

Secondly, numerous regions in both species exhibited significant differences in response to conspecific vocalizations versus non-vocal sounds. These regions were primarily located in the superior temporal gyrus of the temporal lobe and adjacent insula (in marmosets: areas RT, RTM, RM, Tpro, Ipro, PaIL, PaIM, Pir; in humans: areas A1, A4, A5, MBelt, PBelt, LBelt, TA2, 52, RI, PI, STGa, Pir, AVI), as well as anterior temporal areas (STSda and STSva in humans, STR in marmosets). Additionally, humans showed also responses in premotor and inferior frontal regions (55b, 44, 45, IFSp, IFJa, IFJp, OP4), and posterior temporal areas (STSdp, STSvp, TPOJ1 and STV). Interestingly, the cingulate area SCEF in humans and areas 32 and 32v in marmosets also differed in their responses to vocal and non-vocal sounds, particularly later in the testing paradigm. This suggests that both species possess a similar antero-ventral pathway facilitating the identification of diverse vocal information, extending from the superior temporal gyrus to the mid-or anterior cingulate cortex ^1,26^, demonstrating subsequent differentiation in the cingulate cortex after detection occurs in the superior temporal gyrus. These results also indicate that humans possess a posterior-dorsal stream from the posterior temporal cortex to the premotor and prefrontal cortex that is not as pronounced in marmosets.

These shared response profiles suggest some conserved evolutionary pathways for processing conspecific vocalizations in primates, aligning with the model that a network of interconnected voice patches involved in detecting and extracting information from conspecific vocalizations may be preserved across primates ^16–18^. Specifically, the similar response profiles to vocalizations in regions of the medial prefrontal and temporal cortex underscores the importance of these areas in social communication and species-specific vocal recognition in humans and marmosets. This similarity may indicate shared ancestral auditory processing pathways that date back to before the separation of Old and New World primates, which then evolved separately ^31,32^.

However, noteworthy differences emerged between the two species. Firstly, while auditory regions in both species responded to conspecific vocalizations relatively early, some marmoset regions identified non-vocal sounds at content levels of 40-60%, in contrast to all human regions, which required higher levels (70-90%). Furthermore, marmosets exhibited a more gradual response profile for vocalizations compared to the immediate response observed in humans. Human auditory regions discerned vocalizations from white noise at lower mixing levels (20-30% vocalization content), whereas marmosets typically needed higher levels (40-50% vocalization content). This may indicate a higher sensitivity to conspecific vocalizations in humans, potentially reflecting evolutionary adaptations for complex linguistic communications. Therefore, although a similar network was observed between humans and marmosets within the temporal and medial prefrontal cortex, capable of responding earlier to vocal sounds relative to other sounds, the sensitivity of these responses were weaker in marmosets compared to humans.

Behavioral studies corroborate these observations in humans, demonstrating that humans can effectively discriminate voice from non-voice sounds even at very brief sound durations ^33^, and exhibit above-chance vocal emotion recognition from sounds as short as 180 ms ^34^. This suggests an evolutionary refinement of neural circuits that confers a selective advantage for processing communicative sounds. Additionally, the speed with which humans process voice signals is remarkable, as evidenced in the fronto-temporal network. Event-related brain potential studies have highlighted extremely early (150−200 ms) brain responses, indicating rapid discrimination of vocal versus nonvocal sounds ^35^. EEG and MEG studies further support the rapid processing of nonverbal emotional signals from the voice, occurring before 200 ms after voice onset ^36^. These findings, along with major distinctions in voice signal processing from other sounds appearing around 150 ms ^27,28^, reinforce our observations of a highly sensitive network in humans, particularly involving several lateral and medial prefrontal areas and temporal brain regions.

Secondly, a distinct auditory response pattern was observed in marmosets. Some core (A1), belt (R, ML, CL, AL, CM, RTL), and parabelt (CPB, RPB) auditory regions, along with the adjacent TPO area, responded similarly to both vocal and non-vocal sounds or with a slightly higher response to vocalization, but failing to significantly differentiate between the two, showing comparable response levels when comparing these sounds against white noise. In contrast, regions such as the belt area RM and adjacent insular areas AI and Ipro, as well as the lateral temporal Tpro area, exhibited similar responses to both sound types until reaching a threshold of 70-90%, after which the response to vocalizations significantly intensified. This pattern was not observed in humans because even early auditory regions could distinctly differentiate between vocal and non-vocal sounds earlier, responding less to non-vocal sounds.

This broader and less finely tuned sensitivity for vocalizations in these regions of the marmoset suggests an adaptive need for immediate environmental awareness, critical for survival in dense forest habitats. This adaptation suggests that the auditory cortex in primates is tailored to the ecological and social demands of each species ^37^. For instance, recent studies indicate that noise pollution significantly affects marmoset vocal communication, prompting these animals to modify their acoustic communication patterns, which in turn may elevate stress levels and reduce foraging efficiency ^38^. Consequently, the novel environmental sounds included in our study, such as train and aircraft noise, may have been more salient to our marmoset subjects than to humans.

Additionally, both species exhibited regions without significant auditory selectivity. These regions, predominantly located in the peripheral regions of the medial prefrontal cortex in humans (31pd, 33pr, and 8BM) and similar peripheral cortical areas in marmosets (e.g., 29d, 30, 24b), also showed minimal activation during our auditory localizer tasks. This raises intriguing questions about their roles in auditory processing, suggesting that these regions may play minimal roles in vocalization processes or may be involved in other specific auditory tasks.

In conclusion, our study employing comparable paradigms across species provides valuable insights into the similarities and differences in vocalization processing networks between humans and marmosets. Our findings highlighted that both species preferentially process vocalizations, detecting them earlier in an antero-ventral pathway comprising several core, belt, and parabelt auditory areas located in the superior temporal gyrus, as well as adjacent insular areas and anterior temporal areas, along with the cingulate cortex. However, whereas humans also possess a second preferential posterior-dorsal pathway for vocalizations involving the posterior temporal cortex, premotor, and prefrontal cortex, this pathway was not as marked in marmosets. In contrast, the pathway in marmosets involving lateral belt and parabelt posterior temporal areas, as well as the TPO region, processed both vocalizations and non-vocal sounds similarly, showing comparable response levels between the two, which was not observed in humans.

Future studies should consider exploring the temporal dynamics of the processing of the voices of other species, as recent evidence shows that the anterior temporal voice patches of two macaques also possessed a subpopulation of neurons with strong selectivity for human voice ^39^. This could provide further insights into whether these pathways are exclusively dedicated to processing the voices of their conspecifics or are more universally applied across species.

## Methods

### Common Marmoset Subjects

All experimental procedures adhered to the guidelines established by the Canadian Council of Animal Care, under a protocol approved by the Animal Care Committee of the University of Western Ontario (Approval number: 2021-111). The study included nine awake common marmosets (*Callithrix jacchus*): four females (weights ranging from 350g to 450g, aged 48-74 months) and five males (weights ranging from 380g to 464g, aged 44-48 months). These animals participated in both the auditory localizer and dynamic auditory-stimulation paradigm.

To minimize head motion during MRI acquisition, the animals were surgically implanted with an MR-compatible machined PEEK (polyetheretherketone) head post. For comprehensive details of the surgical procedure and the post-operative training period in a mock scanner, please refer to our previous methodological publication by Zanini et al. (2023) ^4040^.

### Human participants

Our study involved nineteen healthy human subjects (9 females; age range: 22-40 years, mean age: 28.7 years). All participants were right-handed, had normal or corrected-to-normal vision, and reported no history of neurological or psychiatric disorders. Participants were fully informed about the experimental procedures and provided written informed consent prior to participation. The experimental protocols were approved by the Ethics Committee at the University of Western Ontario. Human subjects participated in both the auditory localizer and dynamic auditory-stimulation paradigm.

### Stimuli and tasks

The dynamic auditory-stimulation paradigm utilized a range of stimuli including conspecific vocalizations, non-vocal sounds, and pure white noise. To assess auditory response thresholds under diverse conditions, vocalizations and non-vocal sounds were mixed with varying levels of white noise. Specifically, each type of stimulus was presented in a gradient, transitioning from 100% white noise (0% sound) to 0% white noise (100% sound) across 11 equidistant time points. Pure white noise was also presented in a similar stepped format as a control (Figure 10A).

**Figure 10.**
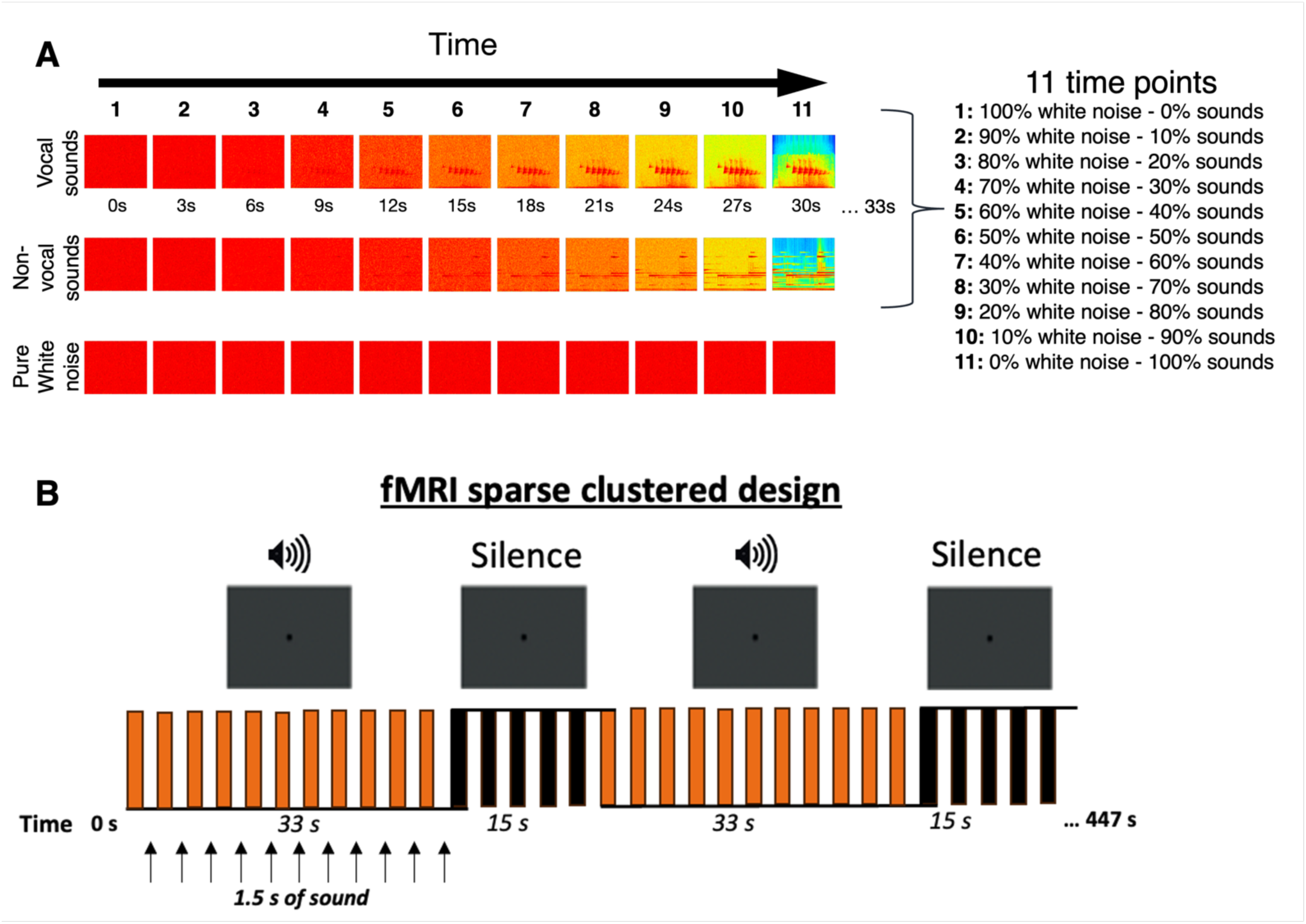
Experimental design and stimulus presentation for the dynamic auditory-stimulation paradigm. **A.** Representation of stimuli used in the paradigm, gradually varying from 100% white noise (0% sound) to 100% sound (0% white noise) over 33 s. Each row indicates a different stimulus condition: conspecific vocalizations, non-vocal sounds, and pure white noise. **B.** Schematic of the fMRI sparse clustered design used in the experiment. The timeline shows the alternation between stimulus blocks (orange bars; each lasting 33 seconds and consisting of eleven 3-second repetition time with 1.5 seconds of acquisition followed by 1.5 seconds of sound presentation) and silence baseline blocks (black bars; each lasting 15 seconds). Each condition was presented three times, resulting in a total of nine stimulus blocks and ten baseline blocks per run.

The gradients were generated using a custom MATLAB script, which linearly adjusted the sound-to-noise ratio at each time point. This script blended vocal or non-vocal sounds with white noise using linear interpolation, gradually decreasing the proportion of white noise while increasing the proportion of the target sound with each step. This meticulous approach ensured that each mixture was equidistant in terms of auditory signal strength from one point to the next, enabling a systematic investigation of auditory sensitivity across a spectrum of sound-to-noise mixtures.

For marmosets, the vocalizations included a variety of conspecific calls: phee, twitter, chirp, and tsik selected from previous recordings ^19^, each trimmed to 1.5 seconds. For humans, the vocalizations consisted of non-speech sounds including laughter, crying, coughing, and other emotive vocalizations provided by Pascal Belin’s group and previously published ^7^, adjusted to the same duration. Twelve different sets of vocalizations were selected for each species, covering all previously mentioned calls for marmosets and non-speech sounds for humans.

Non-vocal sounds included a variety of natural (e.g., sea waves, river water, thunderstorm) and artificial sounds (e.g., bells, helicopter, keyboard), totaling twelve variants from the My Noises App (myNoise, myNoise BVPA). The spectral power of all sounds was normalized using a MATLAB program to maintain consistent auditory levels across stimuli.

Each experimental run featured randomly presented blocks of stimuli alternated with silence periods. Stimulus blocks lasted thirty-three seconds, consisting of 11 three-second intervals (1.5 seconds of acquisition followed by 1.5 seconds of silence where the sounds were presented), interspersed with fifteen-second baseline blocks. Each run included nine stimuli blocks—three each of vocalizations, non-vocal sounds, and white noise—and ten baseline blocks (Figure 10B). To randomize the presentation and minimize order effects, six different sets of stimulus sequences were used, counterbalanced within and across subjects.

For the auditory localizer tasks, we employed the protocol from our previous study on marmosets (see figure 1 in ^15^), which included four continuous 12-second periods of marmoset vocalizations, featuring mainly phee, twitter, tsik, and chatter calls, as well as scrambled vocalizations and a variety of nonvocal sounds comprising natural, man-made, and animal noises. For humans, the protocol mirrored this setup, featuring human non-speech vocalizations, their scrambled versions, and the same non-vocal sounds used for marmosets. The non-speech vocalizations consisted of four 12-second continuous periods using stimuli previously published by Pascal Belin’s group ^7^. The scrambling process involved dividing the signal into multiple frequency bands, scrambling each independently, and then recombining them to mask any linguistic content, following methodologies from our earlier studies ^19,20^. Auditory stimuli were presented in a block design, alternating between stimuli and baseline blocks, with each type of auditory stimulus repeated four times in a randomized order across runs. As previously, the spectral power of all sound files was normalized using a custom program in MATLAB to ensure uniform auditory levels.

### fMRI experimental setup

During scanning sessions, marmosets were placed in a sphinx posture within a custom-designed 3D-printed plastic chair inside a horizontal 9.4 T magnet. Their heads were secured using a head fixation system that clamped onto the surgically implanted head post ^41^. MRI-compatible auditory tubes (S14, Sensimetrics, Gloucester, MA) were inserted bilaterally into the animals’ ear canals, secured with reusable sound-attenuating silicone earplugs and self-adhesive veterinary bandage. The coil housing was then positioned around the head. For further details, see our recent methods paper ^40^.

Inside the scanner, monkeys faced a translucent screen 119 cm away, displaying a small black circle (1.5 cm in diameter) on the center of a gray background projected with an LCSD-projector (Model VLP-FE40, Sony Corporation, Tokyo, Japan) via a back-reflection on a first surface mirror. Auditory stimuli were delivered using Keynote software (version 12.0, Apple Incorporated, CA), synchronized with MRI TTL pulses via a Raspberry Pi (model 3B+, Raspberry Pi Foundation, Cambridge, UK) running a custom-written Python program. Animal monitoring was conducted with an MRI-compatible camera (model 12M-i, MRC Systems GmbH, Heidelberg, Germany), and rewards were given before and after, but not during, scanning sessions.

Human subjects lay in a supine position within the MRI 7T scanner, equipped with MRI-compatible auditory tubes (S15, Sensimetrics, Gloucester, MA). The same black circle with gray background were presented via a rear projection system (Avotech SV-6011, Avotec Incorporated) through a surface mirror affixed to head coil. The auditory stimuli presentation and synchronization process mirrored that of the marmosets, using the same Keynote software and Raspberry Pi setup. Sound levels were carefully adjusted for each participant to maximize comfort, setting the volume just below the threshold of discomfort as reported by each subject.

### fMRI acquisition and parameters

Marmoset and human imaging were conducted at the Center for Functional and Metabolic Mapping at the University of Western Ontario.

Marmosets imaging was performed on a 9.4T/31 cm horizontal bore magnet and a Bruker BioSpec Avance III console equipped with Paravision-7 software (Bruker BioSpin Corp). Data collection utilized a custom-built gradient coil with a 15-cm inner diameter and a maximum gradient strength of 1.5 mT/m/A, alongside an eight-channel receiver ^40,41^.

For both the localizer task and the dynamic auditory stimulation paradigm, eight functional images per task were acquired across four sessions (two sessions per task), utilizing a gradient-echo single-shot echo-planar imaging (EPI) sequence. Imaging parameters included a TR of 3s, acquisition time TA of 1.5s, TE of 15ms, a flip angle of 40°, a field of view of 64×48 mm, a matrix size of 96×128, an isotropic resolution of 0.5 mm³, 42 axial slices, a bandwidth of 400 kHz, and a GRAPPA acceleration factor of 2 (left-right). An additional EPI set for distortion correction featured the opposite phase-encoding direction (right-left). A sparse clustered acquisition paradigm was employed, where each 3-second TR consisted of two phases: the first 1.5 seconds for active RF pulse and gradient acquisition, followed by 1.5 seconds of silence with the gradients turned off but the RF still active. This design ensured that auditory stimuli were presented during periods of scanner silence to minimize noise interference. Each session also included acquiring a T2-weighted structural image with parameters set as follows: TR=7s, TE=52ms, field of view=51.2×51.2 mm, resolution of 0.133×0.133×0.5 mm, 45 axial slices, bandwidth=50 kHz, and a GRAPPA acceleration factor of 2.

Human Imaging was performed on a Siemens Magnetom 7T 68 cm MRI Plus scanner equipped with an AC-84 Mark II gradient coil, an in-house 8-channel parallel transmit, and a 32-channel receive coil ^25^. Five functional runs for the dynamic auditory stimulation paradigm and three for the localizer task were acquired within a single session. Multi-band EPI BOLD sequences were used with the following settings: TR=3s, TE = 20ms, flip angle = 30°, field of view=208×208 mm, matrix size = 104×104, resolution of 2 mm^3^ isotropic, number of slices= 62, GRAPPA acceleration factor: 3 (anterior-posterior), multi-band acceleration factor: 2. Each 3-second TR was structured to include 1.5 seconds of active RF pulse and gradient acquisition, followed by 1.5 seconds of silence with gradients turned off but RF still active, similar to the sparse clustered acquisition employed in marmosets. This setup minimized noise interference during auditory stimulus presentation. Field maps derived from phase and magnitude images were used for artifact correction. Additionally, an MP2RAGE structural scan was acquired for each participant with the following parameters: TR=6s, TE=2.13 ms, TI1 / TI2 = 800 / 2700 ms, field of view=240×240 mm, matrix size= 320×320, resolution of 0.75 mm3 isotropic, number of slices= 45, GRAPPA acceleration factor (anterior posterior): 3.

### fMRI preprocessing

Marmoset fMRI data were preprocessed using AFNI ^42^ and FSL ^43^ software. Initially, raw MRI images were converted to NIfTI format using AFNI’s dcm2nixx and reoriented to match the sphinx position using FSL’s fslswapdim and fslorient functions. Functional images underwent distortion correction with FSL’s topup and applytopup functions, despiking with AFNI’s 3Ddespike to reduce outlier spikes, and time shifts correction using 3dTshift. Images were then registered to the base volume, typically the middle volume of each time series, using 3dvolreg. Spatial smoothing was applied with a 1.5 mm full width at half-maximum (FWHM) Gaussian kernel using 3dmerge AFNI’s function. The mean functional image for each run was calculated and linearly registered to each animal’s respective anatomical image using FSL’s FLIRT after manual skull-stripping using FSL eyes. The resulting transformation matrix was then applied to register all functional volumes to the animal’s anatomical scan, followed by masking to exclude non-brain voxels. Finally, the individual anatomical images were linearly registered to the NIH marmoset brain template ^23^ using Advanced Normalization Tools (ANTs), and the resulting transformation matrix was applied to register functional volumes to the same template. For the localizer task, data were additionally bandpass filtered from 0.01 to 0.1 Hz after smoothing.

Human fMRI data were processed using SPM12 (Wellcome Department of Cognitive Neurology). After converting raw images into NifTI format, functional images were realigned to correct for head movements and underwent slice timing correction. Field map correction was applied using the field map toolbox in SPM to correct for B0 distortion from the magnitude and phase images. The corrected functional images were coregistered with individual MP2RAGE structural scans and normalized to the Montreal Neurological Institute (MNI) standard brain space. Anatomical images were segmented into white matter, gray matter, and cerebrospinal fluid (CSF) partitions and also normalized to the MNI space. Functional images were spatially smoothed using a 4 mm FWHM isotropic Gaussian kernel. For the localizer task, a high-pass filter of 128 seconds was applied to remove low-frequency noise.

### fMRI statistical analysis

#### Auditory localizer task

For each run, we applied a general linear model (GLM) using AFNI’s ‘BLOCK’ convolution to convolve the task timing with the hemodynamic response for each auditory condition. We generated a unique regressor for each of the three conditions to match our 12-second stimulus block experimental design: vocalizations, scrambled vocalizations and non-vocal sounds. These conditions, along with polynomial detrending regressors and motion parameters estimated during realignment for marmosets and humans, were incorporated into the model. This process yielded separate T-value maps for each condition per run, reflecting the neural response to each experimental condition.

These resultant regression coefficient maps from marmosets were then aligned to the NIH marmoset brain atlas template space using the transformation matrices obtained from the registration of anatomical images on the template.

Group-level statistical analyses were conducted using paired t-tests (AFNI’s 3dttest++). Initially, we identified brain regions exhibiting significantly greater activation for each condition – vocalizations, scrambled vocalizations, and non-vocal sounds – compared with the baseline (i.e., vocal > baseline; scrambled vocal > baseline; non-vocal > baseline contrasts), where baseline is defined as the resting state activation recorded during silent intervals between stimuli (15-second baseline blocks). Subsequently, we isolated brain regions preferentially engaged by vocalizations by comparing these to scrambled vocalizations and non-vocal sounds (i.e., vocal > scrambled and vocal > non-vocal contrasts). Z-value maps obtained from these comparisons were thresholded using voxel-level corrections for multiple comparisons (p < 0.01). Significant activations were visualized on fiducial maps using the Connectome Workbench (v1.5.0), employing the NIH marmoset brain template ^23^ for marmosets and the MNI Glasser brain template ^24^ for humans.

#### Dynamic auditory-stimulation paradigm

Based on the voice-sensitive regions identified in our auditory localizer task, we extracted 37 human and 47 marmoset regions of interest (ROIs) from the right hemisphere, using the Paxinos parcellation of the NIH marmoset brain atlas ^23^ for marmosets and the MNI Glasser brain template^24^ for humans. Theses ROIs were categorized by their cortical positions in each species. For marmosets: 6 dorsolateral prefrontal areas, 2 orbital frontal areas, 1 premotor area, 10 medial prefrontal and posterior cingulate areas, 5 lateral, inferior, and ventral temporal areas, 13 auditory areas, 7 insula and other regions in the lateral sulcus, and 3 secondary somatosensory areas. For humans: 6 dorsolateral prefrontal and inferior frontal areas, 1 premotor area, 5 medial prefrontal and posterior cingulate areas, 1 lateral temporal area, 15 auditory areas, 3 insula and other regions in the lateral sulcus, 1 posterior opercular, 1 inferior parietal area, and 4 temporo-parieto-occipital junction areas.

Time courses for each run of each subject were extracted from these predefined ROIs using AFNI’s 3dmaskave function. Data were Z-normalized and segmented according to the specific auditory conditions – white noise, vocalizations, and non-vocal sounds – using condition-specific mask files containing the timing regressor for each condition. This yielded files with 11 data points representing the stimulation period for each condition.

Differences between conditions at each time point were assessed using two-sided paired t-tests, with adjustment for multiple comparisons via False Discovery Rate (FDR) post-hoc correction (p < 0.05), implemented in a custom Matlab script (R022a, The Mathworks). Specifically, we compared the neural responses to vocalizations versus white noise, non-vocal sounds versus white noise, and vocalizations versus non-vocal sounds. This analysis determined the time points at which significant differences between these auditory comparisons emerged for each ROI.

To visually interpret these findings, we generated heat maps and performed hierarchical clustering using Euclidean distance to organize the ROIs into clusters based on the response onsets of significant responses (ranging from 1 indicating very early differences to 11 indicating very late differences).

Finally, to compare auditory responses between species, hierarchical clustering using Euclidean distance was performed combining ROIs from both marmosets and humans. This analysis focused on the presence of differences between conditions in these ROIs, rather than the specific response onsets.

## Acknowledgements

This research was supported by a Discovery grant of the Natural Sciences and Engineering Research Council of Canada to SE. AD was supported by a Canadian Institutes of Health Research postdoctoral fellowship (FRN 193997). Additional support came from the Canada First Research Excellence Fund through BrainsCAN and a Brain Canada Platform Support Grant. We wish to thank Cheryl Vander Tuin, Whitney Froese, Hannah Pettypiece, and Miranda Bellyou for animal preparation and care, Dr. Alex Li and Trevor Szekeres for scanning assistance, and Dr. Kyle Gilbert and Peter Zeman for coil designs.

